# Mercury concentrations in Double—crested Cormorant chicks across Canada

**DOI:** 10.1101/185280

**Authors:** Raphael A. Lavoie, Linda M. Campbell

## Abstract

Mercury (Hg) biomagnifies in aquatic food chains and can reach high concentrations in fish-eating birds. Spatial patterns of Hg have been found in freshwater ecosystems across Canada for many taxa including fish and birds. However, it is often challenging to sample a representative population size of adult birds to monitor concentrations of contaminants over a large spatial scale. Moreover, adult birds can migrate and can show a contaminant profile that may not be representative of local resources. The aims of this study were (1) to determine if there was a spatial pattern of Hg in piscivorous birds, (2) to develop a model to estimate Hg concentrations in breeding adults using chicks as proxy, and (3) to develop predictive equations among non-lethal samples that representative of local resources in adults (blood and growing feathers). Double-crested Cormorant (*Phalacrocorax auritus*) chick growing feathers were sampled at 19 sites across Canada (*n* = 106). Adult tissues (freshly grown feathers; *n* = 8-16 per feather type and blood; *n* = 160) were sampled at five of those locations to establish correlations between age classes and between adult tissues. We found an increase in Hg concentrations with latitude up to 50°N followed by a decrease. There was a decrease in Hg concentrations from west to east, which contradicts previous studies. We found a good correlation of Hg concentrations between adults and chicks and among adult tissues. Our model showed that it is possible to estimate Hg concentrations in adults across Canada using chicks as proxy. Our study shows that chicks can be a suitable proxy for monitoring local mercury concentrations and that they are representative of adults.

*Capsule:* Concentrations of mercury in cormorant chicks are influenced by latitude

## Introduction

Mercury (Hg) biomagnifies in aquatic food chains (Lavoie et al. 2013) and can reach toxic concentrations in upper-trophic-level organisms such as fish-eating birds (Depew et al. 2012, Wolfe et al. 1998). Hg is ubiquitous, but concentrations in wildlife vary spatially depending on the physico-chemistry of the waterbody (e.g., pH) and Hg input in the system (e.g., atmospheric deposition and point sources (Clayden et al. 2013, Kidd et al. 2012)). The trophic structure of a system will also define Hg concentrations in apex predators due to spatial variations in trophic levels (e.g., food chain length (Kidd et al. 2012, Lavoie et al. 2013, Vander Zanden and Fetzer 2007)). Spatial patterns of Hg have been found in freshwater ecosystems across North America for many taxa including fish and bird species and concentrations typically increase towards southeastern Canada and northeastern United States (Depew et al. 2013, Evers et al. 1998, Evers et al. 2003, Scheuhammer et al. 2001, Wente 2004). These patterns are best explained by high deposition of Hg and sulfate, low pH, and surrounding landscapes dominated by forests and/or wetlands (Depew et al. 2013, Driscoll et al. 2007, Evers et al. 2007).

Selecting a representative species to monitor concentrations of Hg over a large spatial scale represents a challenge. Characteristics for good aquatic monitoring species require that they are widespread, abundant, easy to access, easy to capture (and recapture where repeated measurements are necessary), have high territorial fidelity, are high in the food chain (e.g., piscivorous), and have a long lifespan (Evers et al. 1998). Low cost is also an asset in the choice of species used by monitoring agencies. The Double-crested Cormorant (*Phalacrocorax auritus*), an upper-trophic-level piscivorous bird, meet these requirements when nesting on the ground (Dorr et al. 2014, Lavoie et al. 2014, Lavoie et al. 2015). Double-crested cormorants were chosen because chicks are nidicolous (altricial), which means that they require parental care after hatching, such as nourishment, incubation, and protection (Dorr et al. 2014). Cormorant chicks are therefore less prone to escaping when sampled and are available for a long period of time (3-4 weeks for ground nesters and 6-8 weeks for tree or cliff nesters (Dorr et al. 2014)). In addition, cormorant populations are widespread in North America and colonies generally contain many nests (Dorr et al. 2014), making it easier to sample birds without disturbing population sizes. Cormorants show high concentrations compared to other bird species and almost a third of the individuals are at moderate to severe risk to MeHg contamination with many individuals exceeding toxicity benchmarks (Ackerman et al. 2016b). Finally, cormorants embryos are categorized as showing low sensitivity to Hg contamination (Heinz et al. 2009). Yet, field studies in the Great Lakes showed toxic effects of mercury (oxidative stress (Gibson et al. 2014)) and PAHs (DNA mutations (King et al. 2014)) in cormorants. Monitoring resilient species means that if toxicity is found for those conservative species, more sensitive species are likely to show symptoms of toxicity. Considering the reasons enumerated above, we believe that cormorants are a good model species for monitoring and for establishing spatial patterns of contaminants and potentially other chemical tracers such as stable isotopes.

A potential major issue for monitoring studies using birds is that adults of many species migrate and several tissues can show a contaminant profile that may not be entirely representative of local resources of the monitored system (Lavoie et al. 2014). For instance, Hg that was assimilated from food sources at the overwintering area is transferred to the feathers grown during the overwintering period and sequestered as soon as the blood supply to the feathers atrophies. Concentrations measured in such feathers will therefore be representative of the overwintering area even if they were sampled elsewhere (e.g., at the breeding area (Lavoie et al. 2014)). More dynamic tissues (or tissues with fast turnover rate) such as blood and growing feathers will represent a more recent exposure to Hg. Chick tissues can be a good sampling unit because uptake of Hg originates mainly from local food resources (i.e., on breeding grounds) and tissues synthesized locally (e.g., blood or growing feathers) are not influenced by resources from distant ecosystems (Hughes et al. 1997, Stewart et al. 1997). In fish-eating birds, maternal transfer of mercury to eggs or to hatching chicks’ internal tissues have been shown (Ackerman et al. 2011, Ackerman et al. 2016a) and this can bias Hg concentrations because it can come from a mixture of local sources (exogenous reserves) or distant source (endogenous reserves (Becker et al. 1993, Bond and Diamond 2010, Hughes et al. 1997)). However, the effect of maternal transfer is reduced after a few days as chicks grow (biomass dilution and depuration) and concentrations are subsequently defined by local dietary resources (Ackerman et al. 2011). Hg from chick blood is then transferred to growing feathers, which represent short-term local intake of Hg (Bearhop et al. 2000, Hughes et al. 1997). Another reason to use chicks for large-scale monitoring studies is that they are easier to catch than adults, which can drastically reduce time and cost related to sample size requirements.

The objectives of this study was were (1) to determine if there was a spatial pattern of Hg in piscivorous birds from selected breeding sites distributed across Canada, (2) to develop a model to estimate Hg concentrations in breeding adults using chicks as proxy, and (3) to develop predictive equations among non-lethal samples (blood and growing feathers) in adults.

## Materials and methods

### Sample Collection

#### Chicks

Double-crested Cormorant chick feathers were sampled from breeding sites across Canada through associations with other agencies to determine spatial patterns of mercury. The primary feather #1 (hereafter called P1) of chicks was sampled from 19 sites across Canada with four to ten chicks per site for a total sample size of 106 (Figure 1; Table 1) ranging from latitude 42.83 to 54.69°N and from longitude −111.79 to −61.93°W. Chicks were sampled from different nests at each site to keep independence between individuals. Cormorant chicks were caught from accessible nests (e.g., nest on the ground or in low trees). The tip of the growing P1 of the right wing was cut with a pair of scissor. The tip of the feather (which grows first) was sampled to ensure that the same developmental period was sampled among chicks and among colonies. Chick feathers were sampled because uptake of Hg originates almost exclusively from local food sources (i.e., on breeding grounds (Hughes et al. 1997)). In contrast, fully grown feathers in adult birds may contain Hg that was accumulated from resources of distant ecosystems (Lavoie et al. 2014, Lavoie et al. 2015). All feathers were put in paper envelopes until sample preparation. The maximum foraging radius of adult birds during the breeding period is approximately 40 km (Custer and Christine 1992) and a minimum distance of 80 km was therefore kept between colonies to ensure that food sources given by parents to their chicks were distinct among colonies.

**Figure 1.**
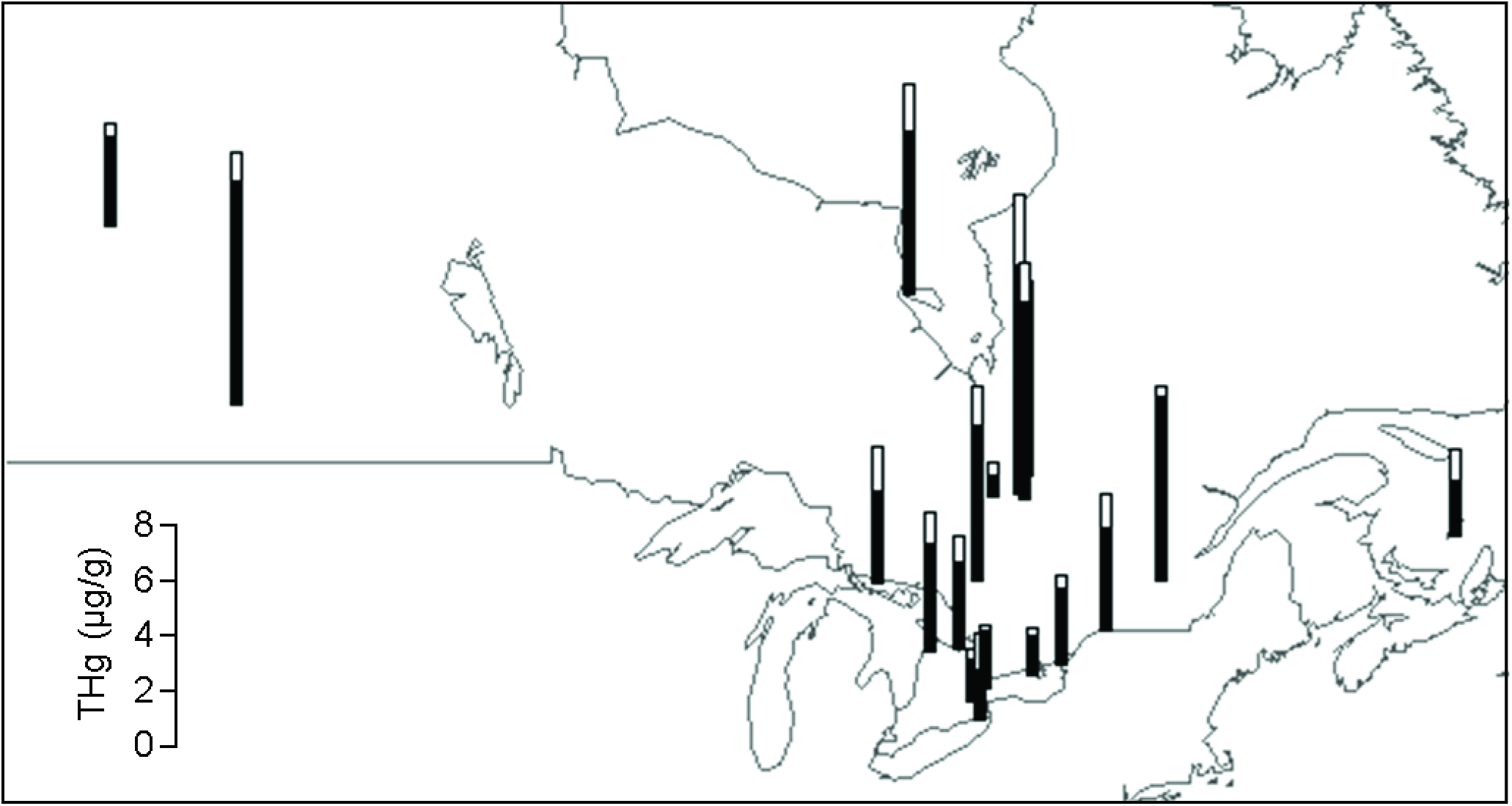
Spatial distribution of Double-crested Cormorant (*Phalacrocorax auritus*) chick mercury (Hg) concentrations across Canada. Black bars represent mean measured Hg concentrations and white bars represent standard deviations (SD).

**Table 1.** Mercury (Hg) concentrations (mean ± standard deviations; SD) in Double-crested Cormorant (*Phalacrocorax auritus*) chick primary 1 feather (P1) from 19 locations across Canada.

| Area | Location | Code | Province | P1 <sup>a</sup> | *n* | Latitude | Longitude | Habitat |
| --- | --- | --- | --- | --- | --- | --- | --- | --- |
| Alberta | Beaver Lake | BICH | AB | 3.3 ± 0.5 | 7 | 54.691 | -111.788 | Lake |
| James Bay | Akimiski Strait | AKI | ON | 5.9 ± 1.7 | 5 | 53.049 | -82.163 | Coastal |
| Lake Erie | Mohawk Island | MOH | ON | 1.8 ± 1.3 | 5 <sup>b</sup> | 42.834 | -79.524 | Lake |
| Lake Huron | Chantry Island | CHAN | ON | 3.9 ± 1.1 | 10 | 44.490 | -81.403 | Lake |
| | Middle Grant Island | MIDD | ON | 3.3 ± 1.7 | 5 | 46.137 | -83.324 | Lake |
| | Nottawasaga Island | NOTT | ON | 3.1 ± 1.0 | 5 | 44.536 | -80.259 | Lake |
| Lake Nipissing | Gull Island | NIP | ON | 5.7 ± 1.4 | 5 <sup>b</sup> | 46.167 | -79.594 | Lake |
| Lake Ontario | Hamilton Harbour | HH | ON | 1.5 ± 0.3 | 10 <sup>b</sup> | 43.284 | -79.794 | Lake |
| | Scoth Bonnet Island | SBI | ON | 1.4 ± 0.3 | 5 <sup>b</sup> | 43.899 | -77.543 | Lake |
| | Snake Island | SNK | ON | 2.7 ± 0.5 | 5 | 44.191 | -76.543 | Lake |
| | Tommy Thompson Park | TTP | ON | 2.1 ± 0.2 | 5 | 43.623 | -79.340 | Lake |
| St. Lawrence River | Bergin Island | BERG | ON | 3.7 ± 1.2 | 5 <sup>b</sup> | 45.014 | -74.860 | River |
| | Lake St-Pierre | LSP | QC | 6.7 ± 0.4 | 4 | 46.198 | -72.849 | Lake |
| Abitibi | Lake Castagnier | CAS | QC | 5.4 ± 1.6 | 5 | 48.729 | -77.764 | Lake |
| | Lake De Montigny | MON | QC | 7.2 ± 1.4 | 6 | 48.144 | -77.907 | Lake |
| | Lake Malartic | MAL | QC | 8.3 ± 2.6 | 5 | 48.256 | -78.104 | Lake |
| | Lake Pelletier | PEL | QC | 0.79 ± 0.4 | 4 | 48.213 | -79.052 | Lake |
| Magdalen Islands | Îlots du Portage | IMAD | QC | 2.0 ± 1.2 | 5 | 47.260 | -61.931 | Coastal |
| Saskatchewan | Reed Lake | REED | SK | 8.1 ± 1.1 | 5 | 50.400 | -107.069 | Lake |

<sup>a</sup> Primary 1 (P1) feathers were sampled at all locations except at Reed Lake where P2 were sampled. <sup>b</sup> Chicks used to pair with adults in the predictive loop model.

#### Adults

Adults were caught at five locations in the Great Lakes area as described in Gibson et al. (2014). Blood (*n* = 160) and growing breast feathers (*n* = 14) were sampled as described in Lavoie et al. (2014, 2015). Briefly, approximately 3 ml of blood was sampled and stored in acid-washed polyethylene vials. Blood samples were stored on icepaks in the field for a maximum of 12 h, centrifuged to keep the cellular fraction (red blood cells) and frozen until analysis. Growing breast feathers were sampled as a result of an artificial molt induced by sampling those same feathers 3-6 weeks earlier. Growing flight feathers (P1: *n* = 8; primaries: *n* = 16; secondaries: *n* = 12; Table 2) were sampled opportunistically throughout the summer by cutting the tip of growing feathers without cutting the blood supply in the sheath. Blood and regrown feathers represent short term exposure of Hg through diet (Bearhop et al. 2000, Monteiro and Furness 2001a, b, Nisbet et al. 2002), although carryover from overwintering locations is possible for birds exposed to high concentrations (Lavoie et al. 2014).

**Table 2.**
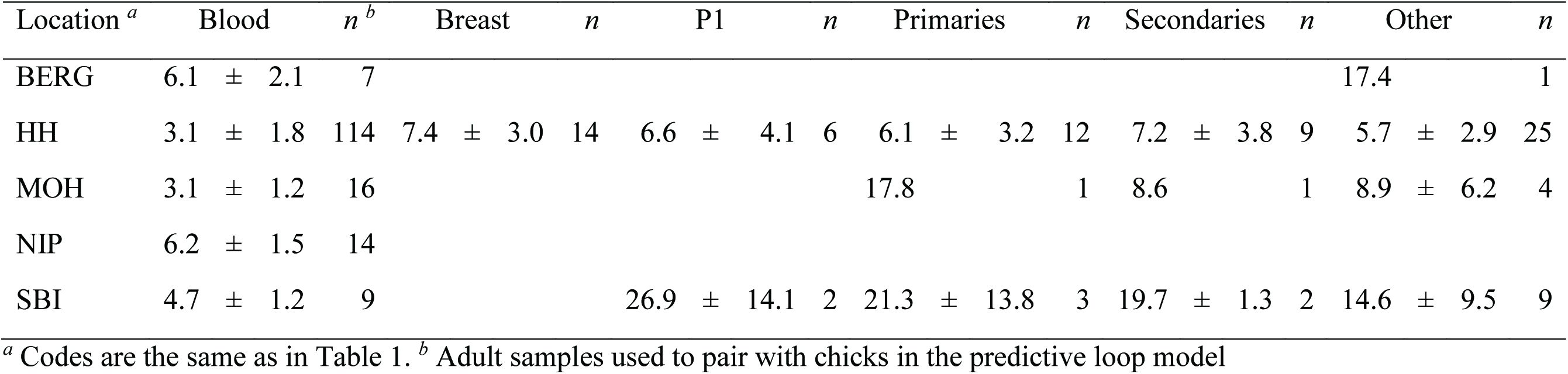
Mercury (Hg) concentrations (mean ± SD) in Double-crested Cormorant (*Phalacrocorax auritus*) adult blood and regrown feathers at 5 locations.

### Sample preparation and mercury analysis

The rachis was cut from all feather samples to avoid any influence of blood supply in the rachis of regrowing feathers (Burger et al. 1992). Each vane of each feather was washed with acetone prior to Hg analysis (Burger and Gochfeld 2000). Red blood cells were freeze-dried prior to analysis (Gibson et al. 2014).

Mercury was analysed in red blood cells and growing feathers as described in Gibson et al. (2014) and in Lavoie et al. (2014). Briefly, samples were digested in a microwave oven with nitric acid, hydrogen peroxide, and deionized water. Total Hg analysis was done by cold vapour atomic fluorescence spectrometry following U.S. EPA method 1631 (U.S. EPA 2002). Certified reference materials from the Canadian National Research Council, Canada (DOLT-4 and DORM-3) and from the Institute for Reference Materials and Measurements, Geel, Belgium (human hair: BCR-397) were analyzed. Mean (±SD) recoveries were 100.6 ± 11.9% (*n* = 33) for DOLT-4, 113.7 ± 14.2% (*n* = 37) for DORM-3 and 96.1 ± 11.2% (*n* = 6) for BCR-397 in adult tissues and 101.2 ± 10.0% (*n* = 12) for DOLT-4, and 110.8 ± 13.2% (*n* = 7) for DORM-3 in chick feathers. Samples THg concentrations are expressed in microgram per gram dry weight (μg/g d.w.) and are assumed to approximate MeHg concentrations in feathers and red blood cells of fish-eating birds (Bond and Diamond 2009, Kim et al. 1996, Lavoie et al. 2010).

### Data analysis

A multiple linear regression (MLR) with chick Hg concentration as the dependent variable and latitude and longitude as independent variables was done. Model selection was done by Akaike Information Criterion corrected for small sample sizes (AICc) to determine the most parsimonious model using the Multi-Model Inference package *MuMIn* (Barton 2013) based on a set of candidate models of main factors and their interaction. When the difference in AICc (ΔAICc) between the best model and the second best model was high (>4) and when Akaike weight (*w;* model probability) of the top model was high, the model output for the model with the lowest AIC was considered to be the most likely model for the dataset. To determine the empirical support of a model over another one, we calculated evidence ratios, which is the Akaike weight of top model over the weight of another model such as the 2^nd^ best model or the null model (i.e. intercept only, (Burnham et al. 2011)).

We developed a model to predict adult Hg concentrations across Canada using concentrations from chicks. Adults (*n* = 160) and chicks (*n* = 30) were caught at five common locations, but could not be paired together because they had no parental link. We therefore generated all possible combinations of regressions between adult blood and chick P1 feathers within each of the five sites (e.g., an adult from site A could only be paired to a chick from site A). Sample sizes used for the model were 5-10 for chicks and 7-114 for adults. We ran a loop of 10,000 iterations of a reduced major axis (RMA) model including all five locations at each iteration. The RMA model was done with the package *lmodel2* (Legendre 2013) with 99 permutations with chick P1 feather concentration as the independent variable and adult blood concentration as the dependent variable. Growing feathers from adults could not be sampled at all five sites and could therefore not be used for the predictive loop model. RMA (Model II) was used as opposed to a simple linear regression (Model I) because both variables were random and were subjected to error. The average slope, intercept, and 2.5-97.5% confidence intervals (CI) of the slope and intercept were used to predict adult blood concentrations. To apply this model to other adult tissues, we ran RMA models between predicted blood concentrations as the independent variable and P1 (*n* = 8), primaries (*n* = 16), secondaries (*n* = 12), breast (*n* = 14), or any growing (*n* = 39) feathers as dependent variables. Hg concentrations were log-transformed for statistical tests to approximate normal distributions.

R software version 3.3.2 (R Core Team 2016) was used for geospatial data treatment, mapping, and statistical analyses.

## Results

### Spatial trends in chicks

Hg concentrations in chick P1 varied from 0.79 ± 0.43 (mean ± SD) μg/g at Lake Pelletier (Quebec; 48.2128°N, −79.0519°W) to 8.3 ± 2.6 μg/g at Lake Malartic (Quebec; 48.2560°N, −78.1042°W; Table 1; Figure 1) and were related to latitude and longitude. A significant polynomial relation with latitude was observed with an increase in Hg concentrations up to approximately 50°N followed by a decrease (*R*^2^ = 0.28, *F*_(1,103)_ = 19.5, *p* < 0.001; Figure 1). We found a decrease in Hg concentrations from west to east, but the model only explained 5% of the variation in Hg (*R*^2^ = 0.05, *F*(1,104) = 4.9, *p* = 0.028). When latitude (polynomial) and longitude were taken together, the best model based on lowest AIC explained 31% of the variation in Hg (multiple linear regression: *F*_(3,102)_ = 15.3, *p* < 0.001):

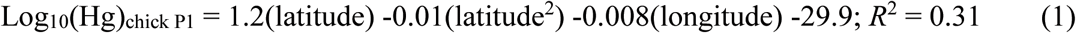

The Akaike weight (i.e., model probability) of the best model was 0.83. The evidence ratio showed that the best model was 4.9 times more likely than the second best model and 15,000 times more likely than the null model.

### Relationship between chicks and adults

The equation of the RMA loop model to predict adult blood Hg concentrations using chick P1 Hg concentrations in the form of *y* = *b* (2.5−97.5% CI) × *x + a* (2.5-97.5% CI; Figure 2), where 2.5−97.5% CI are the lower (2.5%) and upper (97.5%) confidence intervals (CI) was:

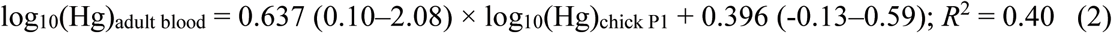

**Figure 2.**
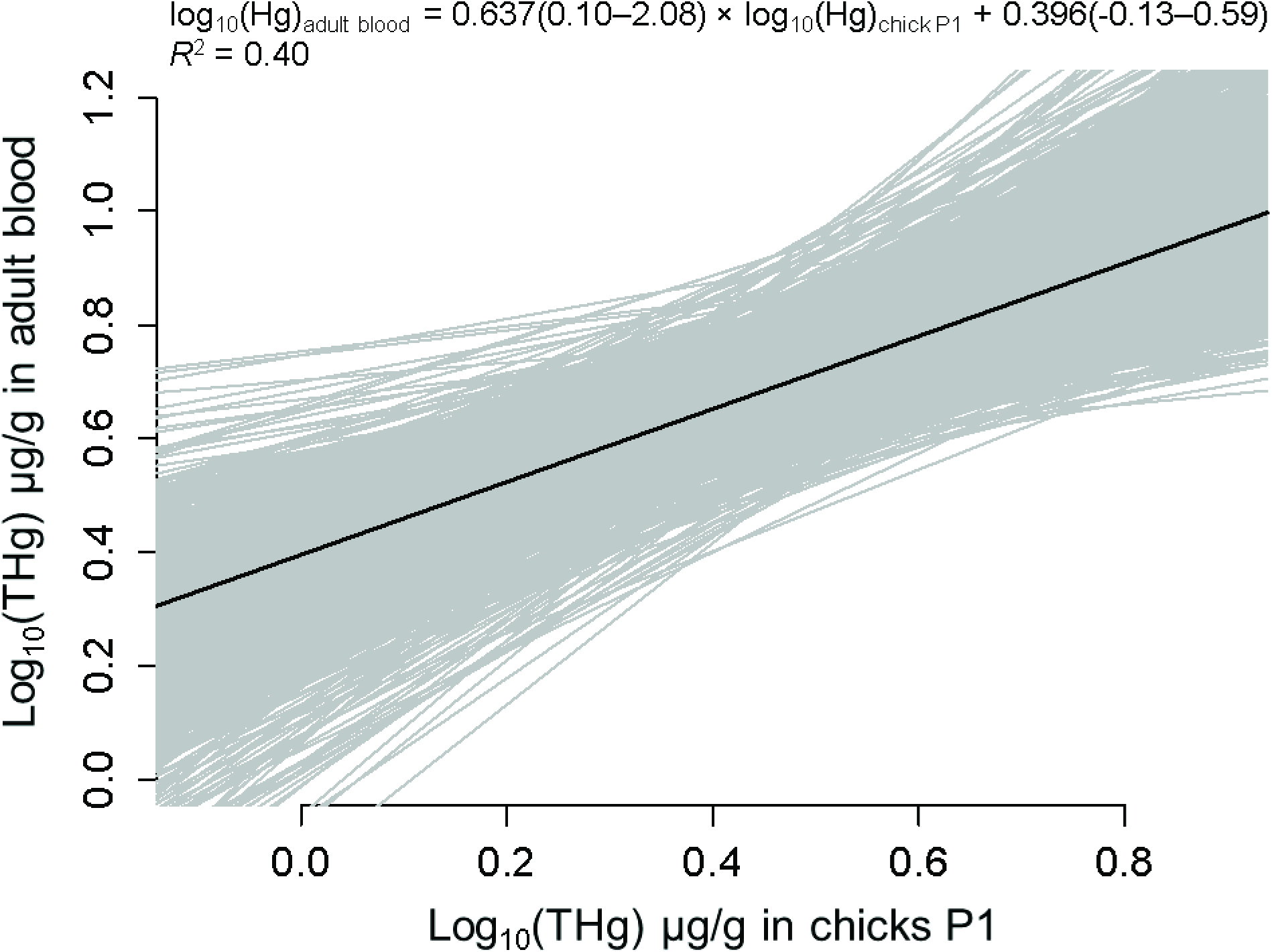
All possible combinations of reduced major axis (RMA) between chicks and adults with average equation shown with solid line and 10,000 iterations shown with grey lines. Average with lower and upper confidence intervals (2.5-97.5%) are shown in the equation.

The CIs showed that the slope was different than 0, but not different than the 1:1 relationship, whereas the intercept was not different than 0.

### Relationships between adult tissues

Hg concentrations in regrown feathers were predicted across Canada using the predicted Hg concentration in adult blood (see equations in Table 3). Predicted adult blood varied from 2.1 ± 0.7 μg/g at Lake Pelletier to 9.5 ± 1.8 μg/g at Lake Malartic (Table 4; Figure 3a). For feathers, concentrations were highest in P1 regrown feathers (3.1 ± 2.2 to 70.8 ± 29.4 μg/g; Figure 3b) and lowest in breast regrown feathers (5.6 ± 1.6 to 19.0 ± 3.0 μg/g) at Lake Pelletier and Lake Malartic, respectively.

**Table 3.**
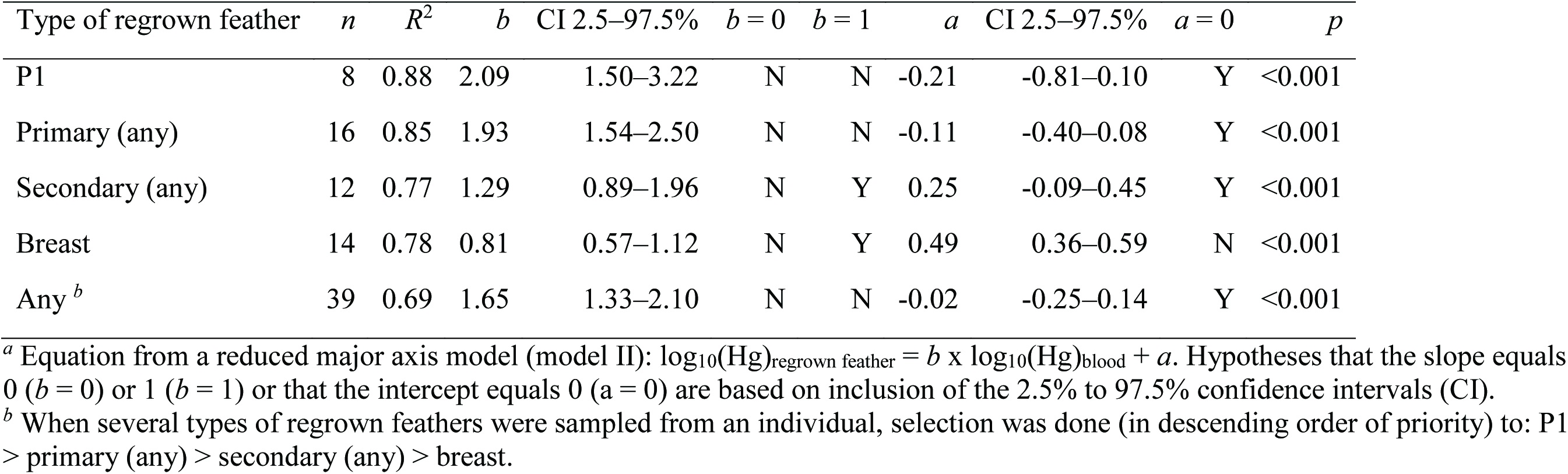
Relationship between log10-transformed mercury (Hg) concentrations in Double-crested Cormorant (*Phalacrocorax auritus*) adult regrown feathers (log10Hgregrown feathers) as dependent variables and adult blood (log10Hgblood) as the independent variable.^*a*^

| Type of regrown feather | *n* | *R*<sup>2</sup> | *b* | CI 2.5–97.5% | *b* = 0 | *b* = 1 | *a* | CI 2.5–97.5% | *a* = 0 | *p* |
| --- | --- | --- | --- | --- | --- | --- | --- | --- | --- | --- |
| P1 | 8 | 0.88 | 2.09 | 1.50–3.22 | N | N | -0.21 | -0.81–0.10 | Y | <0.001 |
| Primary (any) | 16 | 0.85 | 1.93 | 1.54–2.50 | N | N | -0.11 | -0.40–0.08 | Y | <0.001 |
| Secondary (any) | 12 | 0.77 | 1.29 | 0.89–1.96 | N | Y | 0.25 | -0.09–0.45 | Y | <0.001 |
| Breast | 14 | 0.78 | 0.81 | 0.57–1.12 | N | Y | 0.49 | 0.36–0.59 | N | <0.001 |
| Any <sup>b</sup> | 39 | 0.69 | 1.65 | 1.33–2.10 | N | N | -0.02 | -0.25–0.14 | Y | <0.001 |

<sup>a</sup> Equation from a reduced major axis model (model II): log<sub>10</sub>(Hg)<sub>regrown feather</sub> = *b* x log<sub>10</sub>(Hg)<sub>blood</sub> + *a*. Hypotheses that the slope equals 0 (*b* = 0) or 1 (*b* = 1) or that the intercept equals 0 (*a* = 0) are based on inclusion of the 2.5% to 97.5% confidence intervals (CI).
<sup>b</sup> When several types of regrown feathers were sampled from an individual, selection was done (in descending order of priority) to: P1 > primary (any) > secondary (any) > breast.

**Table 4.** Predicted Hg in Double-crested Cormorant (*Phalacrocorax auritus*) adult blood and regrown feathers (mean ± SD)

| Location <sup>a</sup> | Blood <sup>b</sup> | Regrown feathers <sup>c</sup> |  |  |  |  |  |
| --- | --- | --- | --- | --- | --- | --- | --- |
|  |  | Breast | Primaries | P1 | Secondaries | Any |  |
| BICH | 5.3 ± 0.5 | 11.9 ± 0.9 | 19.6 ± 3.5 | 20.3 ± 3.9 | 15.4 ± 1.8 | 15.1 ± 2.3 | |
| AKI | 7.7 ± 1.4 | 16.0 ± 2.3 | 40.8 ± 14.1 | 45.1 ± 16.9 | 25.0 ± 5.8 | 28.1 ± 8.3 | |
| MOH | 3.5 ± 1.5 | 8.4 ± 2.9 | 10.0 ± 9.0 | 10.0 ± 9.9 | 9.3 ± 5.4 | 8.2 ± 6.3 | |
| CHAN | 5.9 ± 1.0 | 13.0 ± 1.7 | 24.6 ± 8.3 | 26.0 ± 9.6 | 17.8 ± 3.9 | 18.2 ± 5.2 | |
| MIDD | 5.2 ± 1.6 | 11.6 ± 3.0 | 20.1 ± 12.7 | 21.1 ± 14.5 | 15.2 ± 6.3 | 15.1 ± 8.1 | |
| NOTT | 5.1 ± 1.1 | 11.5 ± 2.0 | 18.7 ± 7.2 | 19.3 ± 8.0 | 14.7 ± 4.0 | 14.4 ± 4.9 | |
| NIP | 7.5 ± 1.2 | 15.7 ± 2.1 | 38.4 ± 12.0 | 42.3 ± 14.3 | 24.1 ± 5.1 | 26.7 ± 7.2 | |
| HH | 3.2 ± 0.5 | 8.0 ± 1.0 | 7.6 ± 2.1 | 7.3 ± 2.1 | 8.2 ± 1.5 | 6.7 ± 1.6 | |
| SBI | 3.1 ± 0.5 | 7.7 ± 0.9 | 7.0 ± 2.0 | 6.6 ± 2.1 | 7.7 ± 1.5 | 6.2 ± 1.5 | |
| SNK | 4.7 ± 0.5 | 10.8 ± 1.0 | 15.6 ± 3.5 | 15.8 ± 3.8 | 13.2 ± 2.0 | 12.4 ± 2.4 | |
| TTP | 4.0 ± 0.2 | 9.4 ± 0.5 | 11.1 ± 1.3 | 10.9 ± 1.4 | 10.5 ± 0.8 | 9.3 ± 0.9 | |
| BERG | 5.7 ± 1.2 | 12.5 ± 2.2 | 22.9 ± 9.4 | 24.2 ± 10.7 | 16.9 ± 4.7 | 17.1 ± 6.0 | |
| LSP | 8.3 ± 0.3 | 17.1 ± 0.5 | 46.5 ± 3.3 | 51.8 ± 3.9 | 27.6 ± 1.3 | 31.6 ± 1.9 | |
| CAS | 7.2 ± 1.4 | 15.2 ± 2.4 | 35.9 ± 13.6 | 39.3 ± 16.1 | 22.9 ± 5.8 | 25.2 ± 8.1 | |
| MON | 8.7 ± 1.1 | 17.7 ± 1.8 | 51.1 ± 12.4 | 57.5 ± 15.2 | 29.2 ± 4.7 | 34.2 ± 7.1 | |
| MAL | 9.5 ± 1.8 | 19.0 ± 3.0 | 61.8 ± 23.6 | 70.8 ± 29.4 | 32.9 ± 8.3 | 40.0 ± 13.0 | |
| PEL | 2.1 ± 0.7 | 5.6 ± 1.6 | 3.5 ± 2.3 | 3.1 ± 2.2 | 4.7 ± 2.1 | 3.4 ± 1.9 | |
| IMAD | 3.7 ± 1.5 | 8.8 ± 3.0 | 10.8 ± 7.4 | 10.8 ± 7.9 | 9.9 ± 5.0 | 8.8 ± 5.4 | |
| REED | 9.4 ± 0.8 | 18.9 ± 1.3 | 59.5 ± 9.7 | 67.7 ± 12.0 | 32.4 ± 3.6 | 38.9 ± 5.4 | |

<sup>a</sup> Codes are the same as in Table 1. <sup>b</sup> Predicted from chick P1 feathers using Equation 2. <sup>c</sup> Predicted from adult blood using equations from Table 3.

**Figure 3.**
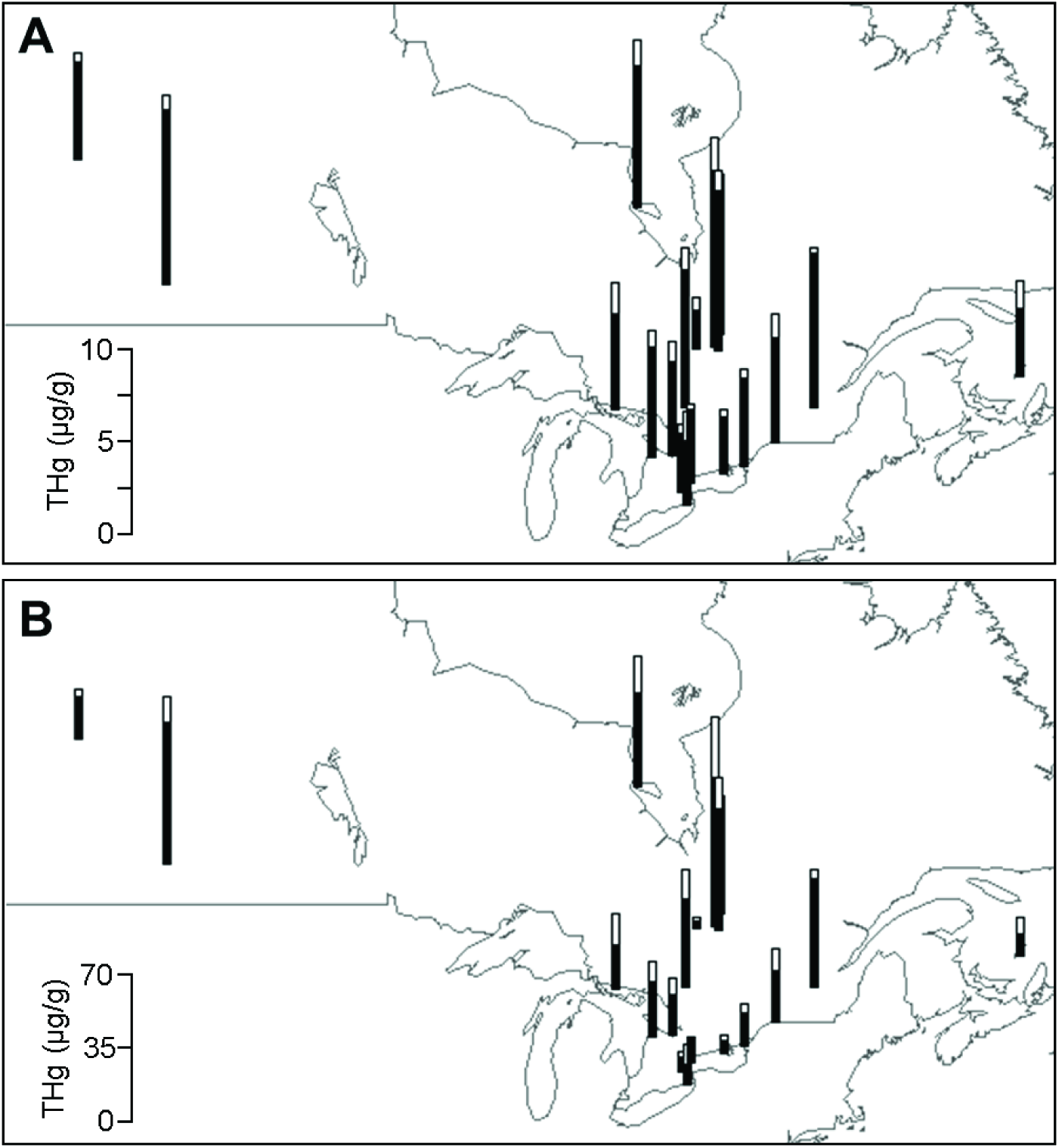
Spatial distribution of Double-crested Cormorant (*Phalacrocorax auritus*) mercury (Hg) concentrations in A) adult blood predicted from chick P1 (Equation 2) and B) adult P1 predicted from adult blood (see equation in Table 3). Black bars represent mean measured Hg concentrations and white bars represent standard deviations (SD).

## Discussion

### Spatial trends

A spatial pattern mainly explained by latitude (polynomial relationship: increase in Hg concentrations up to 50°N followed by a decrease) and longitude (decrease from west to east) was observed. Our work contradicts other studies that found an increase in Hg concentrations from west to east (Depew et al. 2013, Evers et al. 1998, Evers et al. 2003, Scheuhammer et al. 2001, Wente 2004). Several reasons could explain the spatial pattern in this study and the discrepancy between our longitudinal gradient and other studies. First, concentrations of Hg in aquatic organisms are known to vary spatially with Hg inputs. Concentrations of Hg in mosquitoes (Diptera: Culicidae), Largemouth Bass (*Micropterus salmoides*) and Common Loon (*Gavia immer*) were found to be positively related to atmospheric Hg deposition across the U.S. (Hammerschmidt and Fitzgerald 2005, 2006). Hg accumulation in aquatic organisms are therefore greatly influenced by Hg atmospheric loadings and consequently from anthropogenic emission sources (Hammerschmidt and Fitzgerald 2006). Second, Hg concentrations in aquatic biota are greatly influenced by the physico-chemistry of the waterbody. For instance, low pH and high concentrations of ligands, sulfate, and nutrients are central chemical characteristics for promoting Hg methylation in freshwater systems and are associated with increased Hg bioavailability at the water-primary production interface (Dittman and Driscoll 2009, Kidd et al. 2012, Ullrich et al. 2001, Watras et al. 1998). Low water pH has been linked with high Hg concentrations in invertebrates (Chetelat et al. 2011, Clayden et al. 2014) and apex predators such as fish and fish-eating birds (Burgess and Meyer 2008). Methylmercury also binds strongly to dissolved organic carbon (DOC), which can act as a vector of MeHg transport from the catchment area to lakes (Braaten et al. 2014) and can increase concentrations in biota in some cases. Third, the west to east gradient observed in other studies reaches northeastern U.S or southeastern Canada where pH, sulfate and forest type are optimal for Hg bioaccumulation and where Hg hotspots are observed (Depew et al. 2013, Evers et al. 2007, Evers et al. 1998, Evers et al. 2003, Scheuhammer et al. 2001, Wente 2004). Our study is lacking sites in these regions (except for a marine site in Magdalen Islands; 47.2603°N, −61.9311°W with low Hg concentration 2.0 ± 1.2 μg/g) and it is possible that a wider spatial range would lead to a different conclusion. Finally, the trophic structure of a system can also define Hg concentrations in apex predators due to spatial variations in food chain lengths (Kidd et al. 2012, Lavoie et al. 2013, Vander Zanden and Fetzer 2007). We observed that adults in Lake Ontario fed low tropic-level species to their chicks such as Alewife (*Alosa pseudoharengus;* trophic level (TL; mean ± standard error from Fishbase (Froese and Pauly 2016)) = 3.4 ± 0.3) and Round Goby (*Neogobius melanostomus;* TL = 3.3 ± 0.1), while adults in Lake Nipissing fed high trophic level species to their chicks such as Walleye (*Sander vitreus;* TL = 4.5 ± 0.0) and Muskellunge (*Esox masquinongy;* TL = 4.5 ±0.3). Consequently, chicks in Lake Ontario (1.9 ± 0.6 μg/g; *n* = 25) had lower feather Hg concentrations than in Lake Nipissing (5.7 ± 1.4 μg/g; *n* = 5; *t* = −5.8; *p =* 0.003). We believe that spatial patterns are defined by a combination of the factors mentioned above although we cannot identify which ones are the most important ones in this case.

### Relationship between chicks and adults

We found that cormorant chick feathers are a good monitoring tool for large-scale Hg assessments. Chicks of other species have been successfully used for local indicators of Hg pollution or for spatial analyses (Ackerman and Eagles-Smith 2009, Becker et al. 1993, Hughes et al. 1997, Stewart et al. 1997). Cormorants are particularly relevant because they are widespread and abundant across North America (Dorr et al. 2014) and chicks are relatively easy to capture (pers. obs.). In addition, there was a low cost related to the collection of chick samples across Canada because collaborations were made for the sampling (see *Materials and Methods*). Chicks Hg concentrations were found to be representative of those of adults as observed by the linear relationship (Figure 2; Equation 2). Our predictive chick-adult model only explained 40% of the variation, but this is considered as a large effect size according to Cohen (1992). Discrepancies in concentration between chicks and adults can be mainly explained by the sampling design. Chicks were not taken from the same nests as adults and were paired through random iterations. The pairing is therefore at the scale of the colony rather than the nest (i.e, chick-female pairing). In addition, discrepancies could be due to age class differences in bioenergetics and toxicokinetics (e.g., growth dilution (Ackerman et al. 2011, Monteiro and Furness 2001a, b)), types of prey consumed (i.e., adults can bring food to their chicks that may differ than what they eat (Barrett et al. 2007)), and sources of contaminants (i.e., adults may carry contaminants from their migratory habitat (Lavoie et al. 2014); see below). While the match in concentrations between adults and chicks is imperfect, chick values can still be used for spatial patterns assessment of bird colonies because the relative differences among sites should remain regardless of the age class or tissue used. Monitoring cormorant chick feathers could therefore allow the determination of spatial trends and hotspots across North America. In addition, feather sampling of chicks is non-lethal contrary to internal tissues or eggs sampling, which is particularly relevant for endangered species.

Chick feathers is not the optimal choice for monitoring toxicological risks compared to adult blood or eggs (Ackerman et al. 2016b). However, we believe that they can be very useful for the identification of spatial patterns and hotspots for the following reasons. First, chick feathers are relatively easy to obtain and large-scale sampling and large sample sizes can be achieved through collaborations. Second, chicks have limited mobility and are flightless for several weeks and are consequently easier to catch than adults. Third, feather sampling requires little training contrary to blood sampling and collaborations are therefore easier. Finally, shipping feathers across the country or between countries does not require any sample preparation or preservation contrary to blood, eggs or other internal tissues that require proper packing, freezing or drying. This is especially important if samples are retained at the customs for inspection during shipping. Chick feathers are therefore a useful monitoring tool for assessment of large-scale spatial patterns and hotspots.

A potential problem related to the use of chicks is the maternal transfer of Hg. Strong positive relationships have been found between female tissues (e.g., blood) and their eggs or chicks (Ackerman et al. 2016a, Burgess and Meyer 2008, Heinz et al. 2010, Kenow et al. 2015). Because adults are migratory, tissues Hg concentrations may represent a mixture of local resources (exogenous reserves; e.g., breeding area) or distant resources (endogenous reserves; e.g., wintering area (Becker et al. 1993, Hughes et al. 1997, Lavoie et al. 2014)). For species where chicks are covered with down at hatching (e.g., gulls), those feathers are indicative of the mercury in eggs and therefore of the mother (i.e., maternal transfer (Ackerman et al. 2016b)). Females transfer Hg from this mixture of different areas to the eggs, which are then transferred to chicks down. However, cormorants are born naked (no down or feathers) and down growth is only completed after 1-2 weeks (Dorr et al. 2014). The impact of maternal transfer decreases after a few days as chicks grow (biomass dilution and depuration) and local dietary resources subsequently determine concentrations (Ackerman et al. 2011). Hg from blood is then transferred to chicks’ growing feathers, which represent short-term local intake of Hg (Hughes et al. 1997). Cormorant chick primary feathers start to grow by 16-19 days old and they are approximately 2.5 cm long by 21-23 days old (Dorr et al. 2014). Maternal transfer to cormorant chick primary feathers (P1) in our study is therefore unlikely and is believed to represent local dietary and contamination sources.

### Relationships between adult tissues

We found good relationships between blood and different types of growing feathers in adults (*R*^2^= 0.69−0.88), which enables inter-tissues predictions of non-lethal samples. Eagles-Smith et al (2008) found strong relationships among internal tissues (e.g., blood, muscle, liver), but weaker relationships between internal tissues and feathers (head and breast). However, feathers sampled in Eagles-Smith et al (2008) were fully grown feathers that may indicate a mixture of resources provenance. Feathers in our study were regrown feathers, which are good indicators of short-term Hg intake (Bearhop et al. 2000, Monteiro and Furness 2001a, b, Nisbet et al. 2002).

## Acknowledgments

We thank all scientists who contributed to the collection of the chick feather samples: Rod Brook (Ontario Ministry of Natural Resources), Pascale Dombrowski and Jean-Pierre Hamel (*Ministére du Développement durable, Environnement et Lutte contre les changements climatiques)*, Jennifer Doucette (University of Regina), Gail Fraser (York University), Andrea McGregor (University of Alberta), and Jean-François Rail and David Moore (Canadian Wildlife Service (CWS), Environment and Climate Change Canada). We thank Chip Weseloh (CWS), Victoria Donovan, Jason Cox, Laura Gibson, and Kira MacDougall who help in the field and in the lab. This study was supported by a Natural Sciences and Engineering Research Council (NSERC) Canada Research Chair and Discovery funding to L.M.C., an NSERC Alexander Graham Bell CGS-D, a Doctoral research scholarship from the *Fonds Québécois de la Recherche sur la Nature et les Technologies*, an Ontario Graduate Scholarship, and a Senator Frank Carrel Fellowship to R.A.L. This study was also funded by the Wilson Ornithological Society, Bird Studies Canada, and a Graduate Dean’s Travel grant for Doctoral Field Research to R.A.L., and a Canadian Foundation for Innovation and Ontario Research Fund grant to L.M.C.

